# Evidence of non-tandemly repeated rDNAs and their intragenomic heterogeneity in *Rhizophagus irregularis*

**DOI:** 10.1101/205864

**Authors:** Taro Maeda, Yuuki Kobayashi, Hiromu Kameoka, Nao Okuma, Naoya Takeda, Katsushi Yamaguchi, Takahiro Bino, Shuji Shigenobu, Masayoshi Kawaguchi

**Affiliations:** Division of Symbiotic Systems, National Institute for Basic Biology, Japan; School of Science and Technology, Kwansei Gakuin University, Japan; Functional Genomics Facility, National Institute for Basic Biology, Japan; The Graduate University for Advanced Studies [Sokendai], Japan

## Abstract

Arbuscular mycorrhizal fungus (AMF) species are one of the most widespread symbionts of land plants. Our substantially improved reference genome assembly of a model AMF, *Rhizophagus irregularis* DAOM-181602 (total contigs = 210), facilitated discovery of repetitive elements with unusual characteristics. *R. irregularis* has only ten or eleven copies of complete 45S rDNAs, whereas the general eukaryotic genome has tens to thousands of rDNA copies. *R. irregularis* rDNAs are highly heterogeneous and lack a tandem repeat structure. These findings provide evidence for the hypothesis that rDNA heterogeneity depends on the lack of tandem repeat structures. RNA-Seq analysis confirmed that all rDNA variants are actively transcribed. Observed rDNA/rRNA polymorphisms may modulate translation by using different ribosomes depending on biotic and abiotic interactions. The non-tandem repeat structure and intragenomic heterogeneity of AMF rDNA/rRNA may facilitate adaptation to a various environmental condition including the broad host range.

## Introduction

The arbuscular mycorrhizal fungus (AMF) is an ancient fungus with origins at least as old as the early Devonian period^1, 2^. AMF colonizes plant roots and develops highly branched structures called arbuscules in which soil nutrients (phosphate and nitrogen) are efficiently delivered to the host plant^3^. AMF forms symbiotic networks with most land plant species^4, 5^, and the mycelial network formed by various AMF species contributes to plant biodiversity and productivity within the terrestrial ecosystem^6^. The distinctive features of AMF have made it an important model in ecology and evolution^7, 8^; these features include coenocytic mycelia^5^, nutrition exchange with plant, classification as an obligate biotroph^9^, signal crosstalk during mycorrhiza development^9, 10^ and extremely high symbiotic ability^9, 11^.

Recently, multiple genome projects have advanced the understanding of AMF species. Genomic data have been provided for *Rhizophagus irregularis* DAOM-181602^12, 13, 14^, *Gigaspora rosea*^12^*, Rhizophagus clarus*^15^, and other isolates of *R. irregularis*^,14, 16^. These studies revealed potential host-dependent biological pathways^1712^ and candidate genes for plant infection and sexual reproduction ^12, 15, 16^. However, fragmented genome sequences limit the ability to analyze repetitive structures and to distinguish between orthologous and paralogous genes^14^. The first published genome sequence of *R. irregularis* DAOM-181602 (JGI_v1.0)^17^ contained 28,371 scaffolds and an N50 index of 4.2 kb (Supplementary Table 1). The second sequence, by Lin et al. 2014 (Lin14)^13^, contained 30,233 scaffolds with an N50 of 16.4 kb (Supplementary Table 1). Recently published assemblies by Chen et al. 2018 (JGI_v2.0)^14^ contained 1,123 scaffolds with an N50 of 336.4 kb (Supplementary Table 1). The quality of genomic sequence data for other AMF species did not surpass that of DAOM-181602^12, 15^. In contrast, many fungi that are not AMF species contain less than several hundred scaffolds and N50 lengths over 1 Mb^18^. For example, a genomic sequence of an asymbiotic fungus closely related to AMF, *Rhizopus delemar* (GCA000149305.1), was constructed from 83 assemblies with an N50 of 3.1 Mb^19^. Thus, we here present an improved whole-genome sequence of *R. irregularis* DAOM-181602 to facilitate examination of the genomics underlying specific features of AMF species. Taking an advantage of the highly contiguous assembly with little ambiguous regions, we focus on the investigation of the repetitive structures including transposable elements, highly duplicated genes, and rDNA gene copies.

A general eukaryotic genome has tens to thousands of rDNA copies^20^ (Supplementary Fig. 1a), and the sequences of the copies are identical or nearly identical. However, since Sanders et al. (1995)^21^, many studies have indicated intracellular polymorphisms of rDNA (ITS) in various AMF species^22–24^, and the sequencing of isolated nuclei from *Claroideoglomus etunicatum* and *Rhizophagus irregularis* DAOM-181602 suggested sequence variation among the paralogous rDNAs, i.e., intragenomic heterogeneity^13, 25^. This heterogeneity has potentially high impact of studying AMF species, because the rDNA is a fundamental marker of the AMF phylogeny and ecology^8, 26, 27, 28^, and studies have assumed that these rDNAs have no intragenomic sequence variation^29^. Hence, determining the variation degree could cause a reevaluation of the previous understanding of geographic distribution^8^, species identification^28^, and evolutionary processes of AMF. However, the degree of the variation among the 48S rDNA paralogs has been ambiguous because previous studies by Sanger or Illumina sequencing were unable to distinguish each rDNA paralog in a genome. Moreover, the number of rDNA genes in an AMF genome has never been investigated.

The tandem repeat structure (TRS) of the rDNAs is also an attractive topic for evolutionary studies. General organisms require many rDNA copies to make a sufficient amount of rRNA for protein translation^30, 31^. However, in the evolutionary time-scale, multicopy genes reduce in number due to homologous recombination (Supplementary Fig. 1b)^32, 33^ and single-strand annealing (Supplementary Fig. 1c)^34^. To maintain the number of rDNAs, eukaryotes increase the number of copies by unequal sister chromatid recombination (USCR) using the rDNA TRS (Supplementary Fig. 1d,e)^32^. Because this rDNA replacement causes a bottleneck effect in the genome, almost all eukaryotes have homogenous rDNAs in their genomes^20^. This process, termed “concerted evolution,” is an essential system to maintaining eukaryotic protein translation by ribosomes^30^. The heterogeneous rDNAs observed in AMF species implies the collapse of their concerted evolution, and suggest the unique maintenance system of rDNA copy number.

In this study, we built an improved reference genome assembly of *Rhizophagus irregularis* DAOM-181602, which allowed us to discover repetitive elements with unusual characteristics of the AMF genome. We identified unusually small number of rDNA genes in the *R. irregularis* genome. We also found that the rDNA copies are highly heterogeneous and lack a tandem repeat structure.

## Results

### A highly contiguous and complete reference genome of DAOM-181602 generated by PacBio-based *de novo* assembly

We primarily used single-molecule, real-time (SMRT) sequencing technology for sequencing and assembling the *R. irregularis* genome. We generated a 76-fold whole-genome shotgun sequence (11.7 Gb in total) (Supplementary Table 2) from genome DNA isolated from a spore suspension of a commercial strain of *R. irregularis* DAOM-181602 using the PacBio SMRT sequencing platform. A total of 766,159 reads were generated with an average length of 13.1 kb and an N50 length of 18.8 kb (Supplementary Table 2). We assembled these PacBio reads using the HGAP3 program^35^ (149.9 Mb composed of 219 contigs). To detect erroneous base calls, we generated 423 Mb of 101 bases-paired-end Illumina whole-genome sequence data (Supplementary Table 2) and aligned them to the HGAP3 assembly. Through variant calling, we corrected 3,032 single base call errors and 10,841 small indels in the HGAP3 assembly. Nine contigs were almost identical to carrot DNA sequences deposited in the public database (Supplementary Table 3), and these were removed as contaminants derived from a host plant used by the manufacturer. We evaluated the completeness of the final assembly using CEGMA^36^; of the 248 core eukaryotic genes, 244 genes (98.4%) were completely assembled (Table 1 and Supplementary Table 1). Consequently, we obtained a high-quality reference genome assembly of *R. irregularis* DAOM-181602, which is referred to as RIR17.

**Table 1.** Assembly statistics of R. irregularis genome.

|  | RIR17 |
| --- | --- |
| Accession number | BDIQ01000000 |
| Predicted genome size by flow cytometry | 154 Mb |
| Total length of contigs (% of genome) | 149,750,837 bases (97%) |
| # contigs | 210 |
| # N bases | 0 |
| Longest contig (bp) | 5,727,599 |
| Contig N50 (bases) | 2,308,146 |
| L50 | 23 |
| GC % | 27.9% |
| CEGMA completeness for genome contigs | 98.4% |
| # of predicted genes | 41,572 |
| BUSCO completeness for gene models (DB; fungi_odb9) | 94.1% (273/290) |
| Complete single copy | 83.8% (243/290) |
| Complete duplicated | 10.3% (30/290) |
| Fragmented | 3.8% (11/290) |
| Missing | 2.1% (6/290) |

Compared with previous assemblies^13, 14, 17^, RIR17 represents a decrease in assembly fragmentation (1,123 to 210) and an improvement in contiguity using the N50 contig length as a metric (Table 1 and Supplementary Table 1). The total size of the assembly was 9-59 Mb greater than that of previous versions, reaching 97.24% coverage of the whole genome (154 Mb)^17^ (Table 1 and Supplementary Table 1). The new assembly contains no ambiguous bases (N-bases), whereas previous assemblies had 30,115-6,925,426 N-bases (Table 1 and Supplementary Table 1). Approximately 1-7 Mb of sequences from previous assemblies were not contained in RIR17, and JGI_v2.0 has one more conserved gene family than RIR17 (Supplementary Table 1), indicating that a few genomic parts remain to be uncovered by our improvement with continuous sequences. On the other hand, RIR17 was aligned with 95-99.2% of previous assemblies (Supplementary Table 4), suggesting that RIR17 covers the majority of the previously sequenced areas with high sequence contiguity. Moreover, RIR17 contained 8-47 Mb of regions unassigned in previous genomes (Supplementary Table 4). These regions are newly revealed by our improvement.

RIR17 contains a greater extent of repetitive regions than JGI_v2.0. The RepeatModeler^37^ and RepeatMasker^37^ pipeline identified 64.4 Mb (43.03%) of RIR17 as repetitive regions (Supplementary Table 5). These regions total 18.9 Mb more than those of JGI_v2.0 (Supplementary Table 5). Previous fosmid sequences predicted that DAOM-181602 contains ∼55 Mb of repetitive regions^17^, suggesting that RIR17 covers the majority of the repetitive regions of DAOM-181602.

We confirmed a unique repeat profile in the AMF genome. The majority of the interspersed repeats (62.83%) could not be categorized with known repeat classes (Supplementary Table 5), indicating that the AMF genome accumulated novel classes of interspersed repeats. Moreover, DAOM-181602 lacks short interspersed nuclear elements (SINEs), which are abundant in closely related fungi (Supplementary Table 5). Several types of SINEs proliferate using transposases on long interspersed nuclear elements (LINEs)^38^. Although the AMF has 23 LINEs containing the transposase gene (Supplementary Table 6), SINEs have never been found in previous genomes^13, 14, 17^ or RIR17. DAOM-181602 may have a system to suppress the invasion and proliferation of SINEs (e.g., a high number of very active Argonaute proteins, as predicted by Tisserant et al.^17^).

### New gene annotation for DAOM-181602 details gene family expansion in AMF

Using the RIR17 assembly together with strand-specific RNA-Seq data (“Rir_RNA_SS” in Supplementary Table 2), we built a set of 41,572 gene models (Supplementary Tables 6 and 7). Of the genes predicted, 27,859 (67.0%) had either RNA-Seq expression support, homology support or protein motif support (Supplementary Tables 6 and 7). The gene models having any support were submitted to the DDBJ as standard genes and were used in downstream analyses. The models having no support were assigned as “PROVISIONAL” gene models (Supplementary Tables 6 and 7). Using Orthofinder with previous genomic gene sets indicated that our gene models cover the majority of previously provided genes (Supplementary Fig. 2). Although new models showed more coverage of “Benchmarking Universal Single-Copy Orthologs” (BUSCOs)^39^ (Supplementary Table 7) than JGI_v1.0 and Lin14, their gene completeness was slightly lower than that of JGI v2.0 (9 BUSCO families overlooked, Supplementary Table 7), indicating the advantage of using the JGI annotation pipeline to discuss the gene variety in DAOM-181602 (Chen et al 2018^14^). However, we considered our model set suitable for the analysis of the repetitive region and highly paralogous genes because our model is based on highly continuous assemblies, and the number of genes on repetitive regions was increased to 2,349-12,559 genes from the number in JGI_2.0 (Supplementary Table 8).

*R. irregularis* has one of the largest numbers of genes in fungi (Supplementary Fig. 3). Our ortholog analyses indicates that the gene number inflation was caused by lineage-specific expansions of gene families and not by whole-genome duplications. An Orthofinder analysis of nine fungal genomes and two animal data sets (Supplementary Table 9) showed that many of the single-copy genes in other fungi were also single copies in RIR17 (216/239 families, Supplementary Table 10), negating the possibility of whole-genome duplication in *R. irregularis*. The large number of “species-specific single-copy” (SSSC) genes in DAOM-181602 (10,354 genes, Fig. 1a, Supplementary Table 10) suggests that the AMF genes inflated by new gene constructions through gene fusion and mutation accumulation. Moreover, several common gene families in Opisthokonta also contributed to gene inflation; the *R. irregularis* lineage had 92 rapidly expanded (RE) families, containing 8,952 genes (Fig. 1a, d-e, Supplementary Tables 10 and 11), suggesting that *R. irregularis* has also acquired many genes by the duplication of particular gene families.

**Fig. 1.**
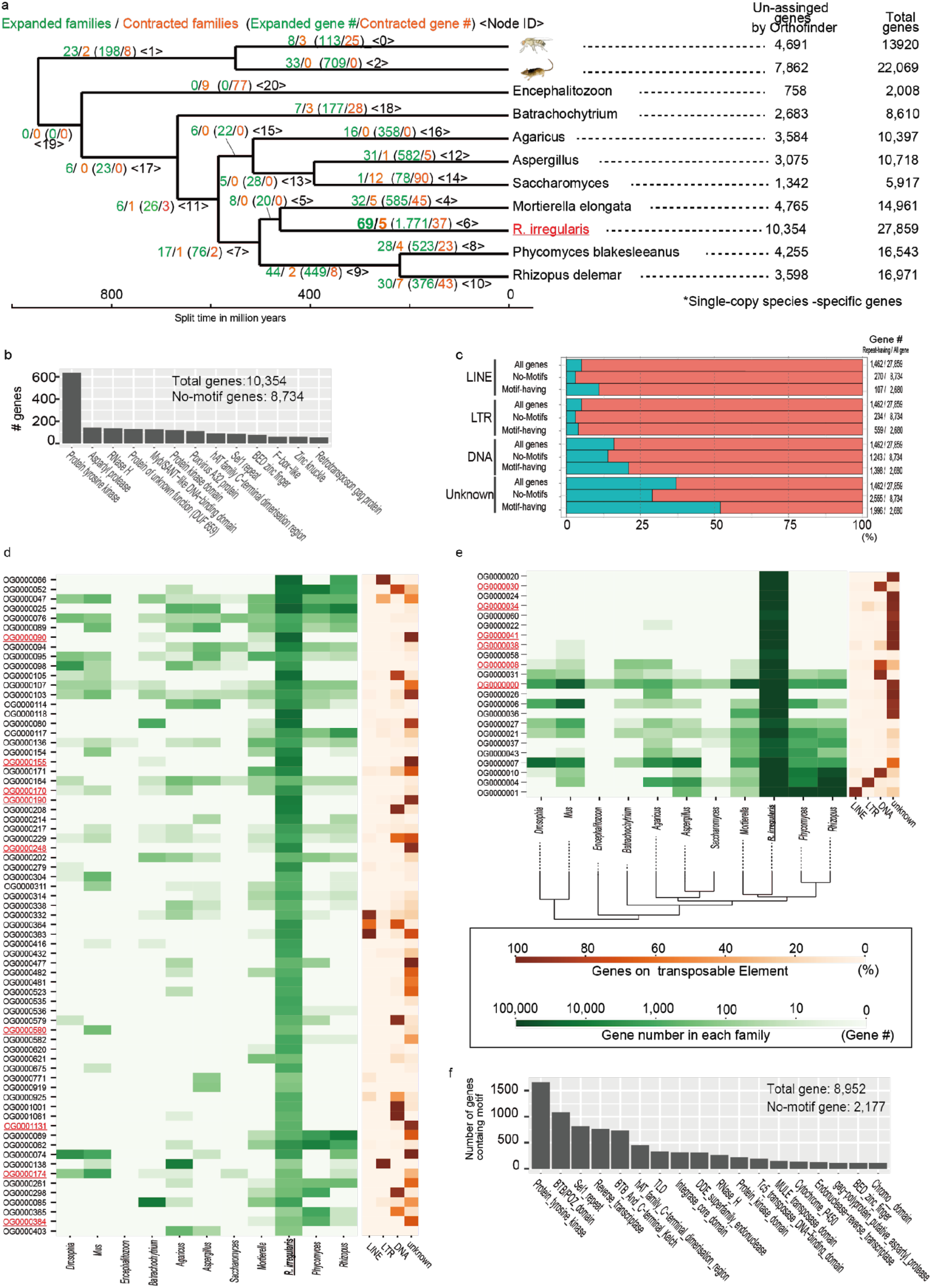
Gene inflation in *R. irregularis*. **a. Rapidly expanded/contracted ortholog groups based on CAFE analysis.** Total gene number of analyzed species and unassigned genes by Orthofinder analysis (species-specific single-copy genes) are described on the right side of the tree. **b. The number of *R. irregularis-*specific single-copy genes having protein motifs**. Minor motifs (<50 genes) were omitted from the Fig. (raw-data; Supplementary Table 12). **c. The proportion of genes among the SSSC genes having each repeat element. d. Sixty-nine rapidly expanded orthologous groups (OGs).** Green heat map shows the number of genes in each OG. Orange heat map indicates the proportion of genes with each repeat element. The OGs containing the “protein tyrosine kinase” domain are marked in red. **e. Rapidly expanded OGs based on z-score analysis.** The colors have the same meaning as in 1d. **f. The number of rapidly expanded ortholog genes having protein motifs.** Minor motifs (<100 genes) are omitted from the Fig. (raw-data; Supplementary Table 12).

The motif annotation indicates that inflated genes may contribute to signaling pathways of AMF species. Our Pfam search annotated 1,620 SSSC genes and 6,755 RE genes (Fig. 1b, f, Supplementary Tables 6 and 10). The most frequently observed motif was “Protein tyrosine kinase; PF07714” (Fig. 1b, f, Supplementary Table 12), which is often found in signaling proteins in multicellular organisms^40^, consistently with previous genome studies^14^. Other signal-related motifs (e.g., Sel1 repeat and BED zinc finger) were also found in the inflated genes (Fig. 1b, f, Supplementary Table. 12). AMF has developed a unique signal pathway for symbiosis (e.g., establishments of symbiosis with pathways using *sis1* and strigolactones^41^). This inflation of signaling-related genes may have led the development of a complex signaling pathway in AMF.

We then investigated the contribution of the transposable elements (TEs) to gene inflation based on the overlapping of highly paralogous genes and the TEs. Previous studies hypothesized that the gene inflation in *R. irregularis* relates with the expansion of TEs^14^. Our analysis showed that in several RE families (e.g., OG0000090 and OG0000020), over 90% of the genes were located with TEs (Fig. 1d,e, Supplementary Table 13), suggesting that TEs accelerated the gene expansion in these families. However, some of the families had no correspondence with TEs (e.g., “OG0000025”, and “OG0000058” in Fig. 1c, e). In SSSC genes, TEs were slightly more frequently found in SSSCs with motifs than in all gene sets but were less frequently found in SSSCs without motifs (Fig. 1c). This detailed analysis supports the contribution of TEs to gene inflation in several gene families but also clarified that several families show TE-independent expansion. Although more genome data for AMF species and sister groups are required to reveal the gene expansion process and its contribution to AM symbiosis, our data provide a fundamental dataset to reveal the evolution of gene redundancy in AMF species.

### Losing conserved fungal genes

Previous AMF studies suggested the loss of several categories of genes by symbiosis with plant^12, 13, 17^. Our RIR17 genome assembly confirmed the loss of genes involved in the degradation of plant cell walls such as cellobiohydrolases (GH6 and GH7 in the CAZy database), polysaccharide lyases (PL1 and PL4), proteins with cellulose-binding motif 1 (CBM1), and lytic polysaccharide monooxygenases (Supplementary Tables 6 and 14) and nutritional biosynthetic genes, including fatty acid synthase (FAS) and the thiamine biosynthetic pathway (Supplementary Table 15). Given that fatty acids and thiamine are essential nutrients for fungi^42, 43^, *R. irregularis* should take up those essential nutrients from a host plant without digestion of the plant cell wall. Several recent papers have described the transport of lipids from plants to AMF^44–46^.

### *R. irregularis* has an exceptionally low rDNA copy number among eukaryotes

The general eukaryotic genome has tens to thousands of rDNA copies^20^ (Supplementary Fig. 1a). However, the RIR17 genome assembly contained only ten copies of the complete 45S rDNA cluster, which was composed of 18S rRNA, ITS1, 5.8S rRNA, ITS2, and 28S rDNAs (Fig. 2a, Supplementary Table 16). To confirm that no rDNA clusters were overlooked, we also estimated the rDNA copy number based on the read depth of coverage. Mapping the Illumina reads of the genomic sequences (“Rir_DNA_PE180”) onto the selected reference sequences indicated that the coverage depth of the consensus rDNA was 8-11 times deeper than the average coverage depth of the single-copy genes (Fig. 2b, Supplementary Table 17), the number of rDNA copies is approximately 10, and the RIR17 assembly covers almost all of the rDNA copies. This AMF rDNA copy number is the lowest among eukaryotes^47^ other than pneumonia-causing *Pneumocystis* (one rDNA)^48^ and malaria-causing *Plasmodium* (seven rDNAs)^49^.

**Fig. 2.**
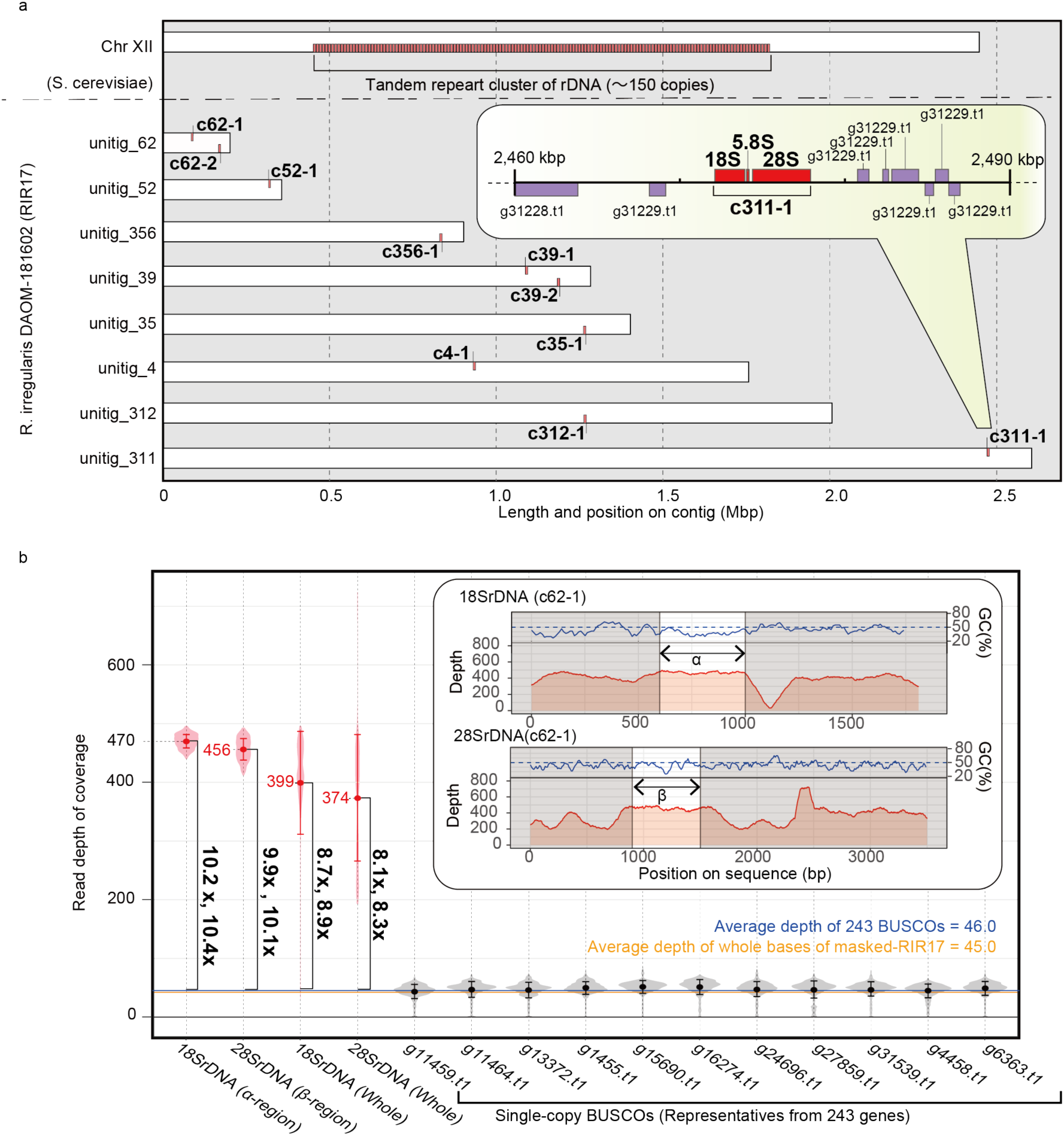
Physical maps of rDNA structures and copy numbers in RIR17. **a. Distribution of *R. irregularis* rDNA units in the genome.** Each 48S rDNA unit is represented as a red box. For comparison, rDNA clusters on *Saccharomyces cerevisiae* chromosome XII are shown^83, 84^. Inset is a magnified view of a 48S rDNA unit (c311-1) with nearby protein-encoding genes (purple boxes). Genes encoded by the plus-strand genome are depicted on the top side, and those encoded by the minus strand are shown on the bottom side. **b. Copy number of rDNA in DAOM-181602 based on the read depth of coverage.** Averages of the “read depth of coverage” are represented as dots and with italic labels. Error bars and violin plots show standard deviations and normalized coverage distribution. The depths of rDNA regions are marked in red. For comparison, the data from representative single-copy BUSCO genes on RIR17 are shown in black. The mean depth of means from 243 BUSCOs is marked with a horizontal blue line, and the mean depth of all RIR17 bases is marked with an orange line. The changes in the depth of rDNA regions are in vertical bold labels and square brackets. rDNA regions adapted for the copy number estimation (α- and β-regions) are marked in the inset with the depth of coverage and the GC content of each sequence position.

This low copy number suggests a unique ribosome synthesis system in AMF. The rDNA copy number has relevance for the efficiency of translation because multiple rDNAs are required to synthesize sufficient rRNA. For instance, an experimental decrease in rDNA copy number in yeast (approximately 150 rDNAs in wild type) resulted in no isolated strain having <20 copies, which is considered the minimum number to allow yeast growth^30^. The doubling time of yeast with 20 rDNA copies (TAK300) was 20% longer than that of the wild type^30^. In DAOM-181602, successive cultivation in an infected state with a plant has been widely observed, suggesting that this exceptionally small rDNA copy number is enough to support growth. The multinucleate feature of AMF would increase the rDNA copy number per cell and thereby perhaps supply enough rRNA to support growth. A similar trend in rDNA reduction is observed in the organellar DNA (e.g., mitochondria and plastids)^50^. Revealing the details of translation in AMF requires a future tracking study of the rRNA production and degradation process in AMF. Elucidation of the mechanism to produce mass rRNAs from a few rDNAs may contribute the understanding of not only AMF evolution but also other polynuclear cells (e.g., striated muscle and ulvophyceaen green algae) and symbiont-derived organelles.

### *R. irregularis* rDNAs are heterogeneous and completely lack a TRS

Interestingly, none of the RIR17 rDNAs form a TRS, in contrast to most eukaryotic rDNAs, which comprise tens to hundreds of tandemly repeated units^20^. Most of the rDNA clusters in RIR17 were placed on different contigs; a single copy of rDNA was found in “unitig_311”, “_312”, “_35”, “_356”, “_4”, and “_52”, and two copies were found in “unitig_39” and “_62” (Fig. 2b, Supplementary Table 16). In the cases where two rDNA clusters were found, the two copies resided apart from each other and did not form a tandem repeat; the distances between the clusters were over 70 kb (76,187 bases in unitig_62 and 89,986 bases in unitig_39, Fig. 2b, Supplementary Table 16), the internal regions contained 31 and 42 protein-coding genes, respectively, and the two clusters were located on reverse strands from each other (Fig. 2a, Supplementary Table 16). Since all rDNA copies are located over 28 kb away from the edge of each contig (Fig. 2a, Supplementary Table 16), the lack of TRSs is unlikely to be an artifact derived from an assembly problem often caused by highly repetitive sequences.

The lack of tandem rDNA structure was also supported by mapping our PacBio reads to RIR17 and searching for rDNA on JGI_v2.0 assemblies. BWA-MEM^51^ mapping showed multiple PacBio reads across the 5′ non-coding region, 48S rDNA and 3′ non-coding regions of each rDNA contig (Fig. 3a, Supplementary Fig. 4). Because our PacBio analysis directly sequenced the DNA molecules in AMF, this syntenic structure is not due to chimeric fragments from DNA amplification but reflects the natural sequence. The 5′ and 3′ non-coding regions of each rDNA have sequences that are not similar other than the highly similar 5′ regions on c62-1 and c62-2 (Fig. 3b and Supplementary Fig. 4), negating the possibility of mapping confusion due to sequence similarity. We reproducibly obtained the PacBio reads passing the rDNA regions from our three PacBio datasets, which had been constructed from different spore suspensions. Furthermore, our rDNA searching by RNAmmer detected a non-tandem 48S rDNA region from three JGI_v2.0 scaffolds (Fig. 3c, Supplementary Fig. 5 and Supplementary Table 18). Although the seven rDNAs cannot be reconstructed from JGI_v2.0, two partial rDNA sequences on JGI_v2.0 had corresponding down- or upstream sequences that matched our RIR17 rDNAs (Supplementary Fig. 5, and Supplementary Table 18), indicating that our assembly around the rDNA genes is consistent with previous assemblies.

**Fig. 3.**
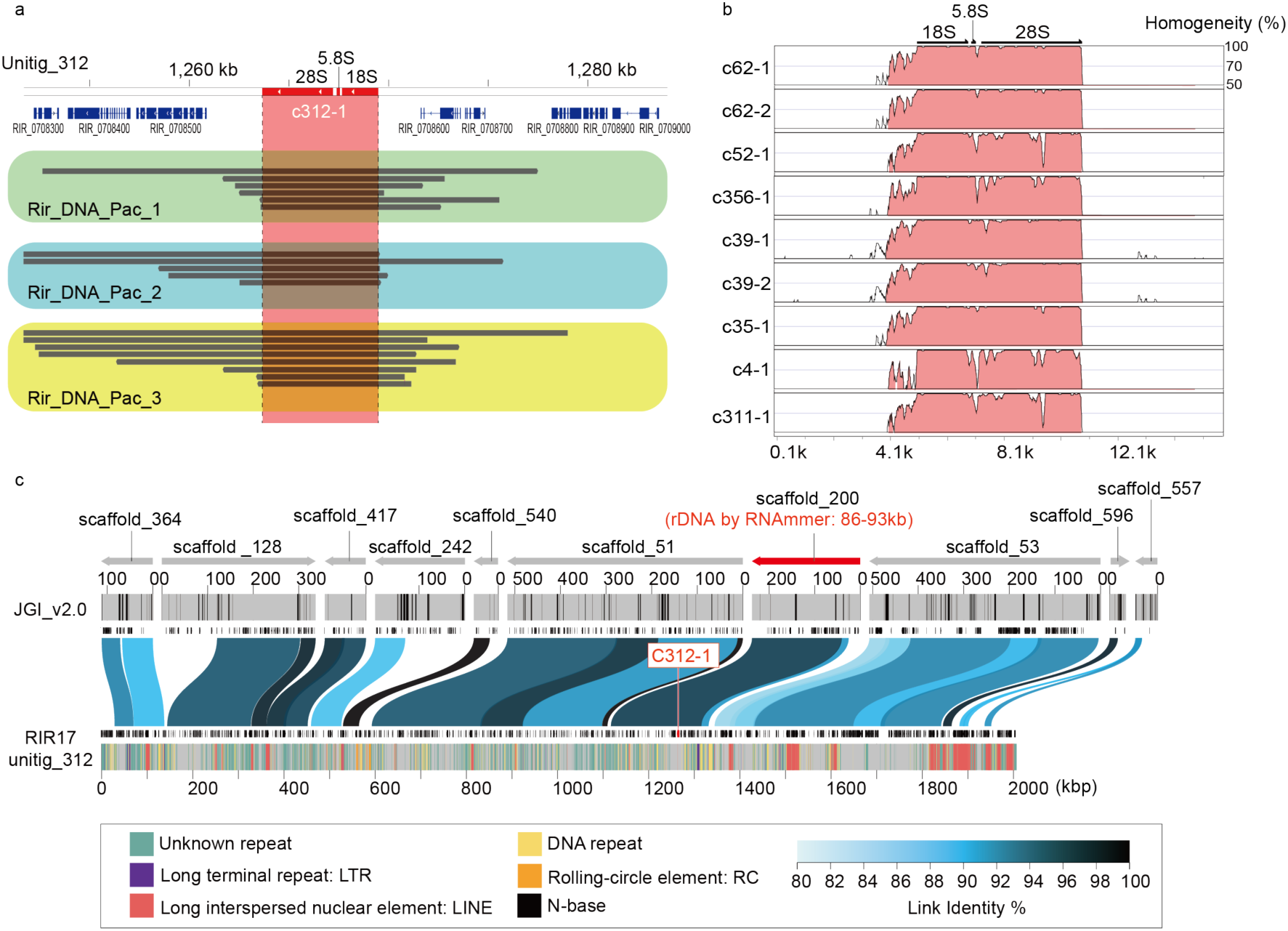
Evidence for the lacking of tandem repeat structures of rDNA. **a. Mapped PacBio read for the rDNA regions on unitig_312 contig in RIR17.** The top bar and tick marks indicate sequence positions on the contig. The rDNA region (c312-1) is indicated in red. Blue boxes show the predicted protein-coding genes. The mapped read was indicated black bar, and reads from different DNA samples and libraries (Supplementary Table 2) boxed with green, light-blue and yellow colors, in each. Mapped reads for the other rDNA regions were summarized in Supplementary Fig. 4. **b. Sequence similarity of c312-1 rDNAs with other rDNA regions on RIR17.** The 5 kb upstream and downstream sequences of each rDNA region are separated from each contig. Alignment and similarity were calculated with mVISTA^85^. Red color shows the sequence regions with similarity over the threshold (>70% similarity for 100b). **c. Positions and identities of JGI_v2.0 scaffold aligned against unitig_312 contigs of RIR17.** The top area indicates aligned scaffolds and their strands. A scaffold containing the predicted rDNA gene is marked in red. The positions of N-base on JGI_v2.0 are marked with black bars in the next line. Predicted protein-coding genes from Chen et al^14^. are indicated with the next black boxes. Aligned positions and their similarity are marked with blue or black bands on the next line. The area below the black boxes show the predicted genes in the present study. Repetitive regions are marked with colored lines on the bottom band. Types of repetitive elements and the legend of similarity coloration are indicated in the bottom box.

We then examined polymorphism among the 45S rDNA clusters on RIR17. rDNA heterogeneity has been reported in various AMF species, including DAOM-181602^13, 17, 25, 29^. However, the distribution and degree of the variation among the rDNA paralogs were unclear. Pairwise comparisons of the ten rDNA copies detected 27.3 indels and 106.1 sequence variants with 98.18% identity on average (Supplementary Tables 19 and 20), whereas the sequences of rDNA clusters at c311-1 and c52-1 were identical. Polymorphisms were distributed unevenly throughout the rDNA; percent identities were 99.91% in 18S rDNA, 97.93% in 28S rDNA, 96.65% in 5.8S rDNA, 93.45% in ITS1, and 90.28% in ITS2 (Fig. 4, Supplementary Tables 19 and 20). The number of polymorphic sites in *R. irregularis* rDNAs reached 4.07 positions per 100 bases, much higher than in other fungi, which have polymorphic sites at 0.04-1.97 positions per 100 bases (Table 2). The rDNA polymorphisms observed in RIR17 covered most of the polymorphisms previously reported in this species (Fig. 5), providing incentive to review the molecular ecology of AMF. The degree of intragenomic variation was not high enough to disrupt species-level identification but was sufficient to cause erroneous identification of *R. irregularis* strains (Fig. 5). These findings pose a caution that previous studies on geographic distribution^8^, species identification^28^, and evolutionary processes of AMF assuming rDNA homogeneity require reevaluation considering the high-level intra-genomic heterogeneity of rDNA sequences in AMFs.

**Fig. 4.**
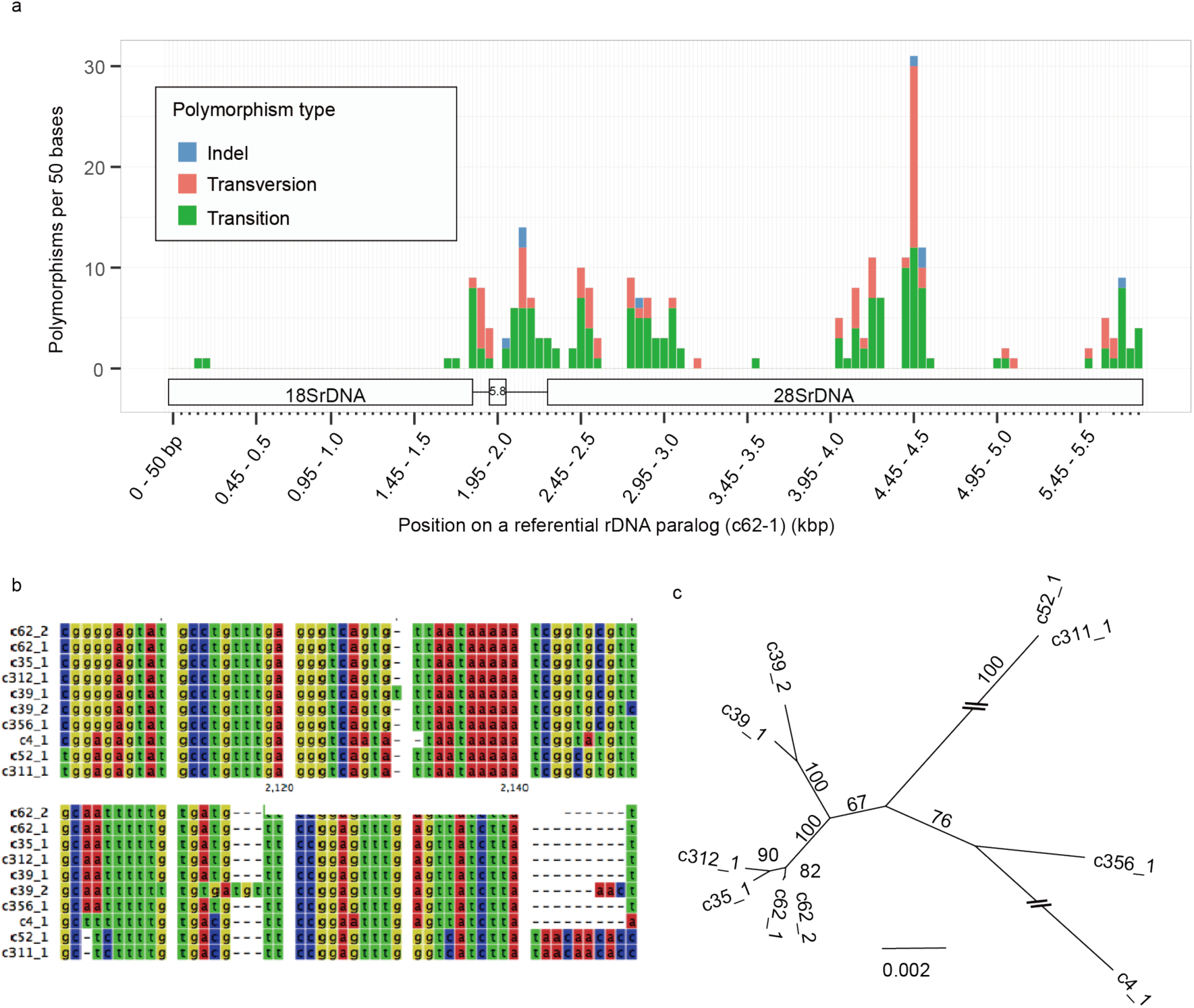
Polymorphisms of 48S rDNA paralogs in RIR17. **a. The distribution of rDNA sequence variants within the 48S rDNA of RIR17**. The position and types of polymorphisms were called based on the paralog c62-1. **b. Alignment of a heterogeneous region among the 48S rDNA paralogs**. Partial sequences of MAFFT-aligned 48S rDNAs (corresponding to 2,049-2,136 bases positions on c62-1). **c. Neighbor-joining tree for phylogenetic relationships among the ten rDNA paralogs based on 5,847 aligned positions.** Bootstrap values are described at each node.

**Fig. 5.**
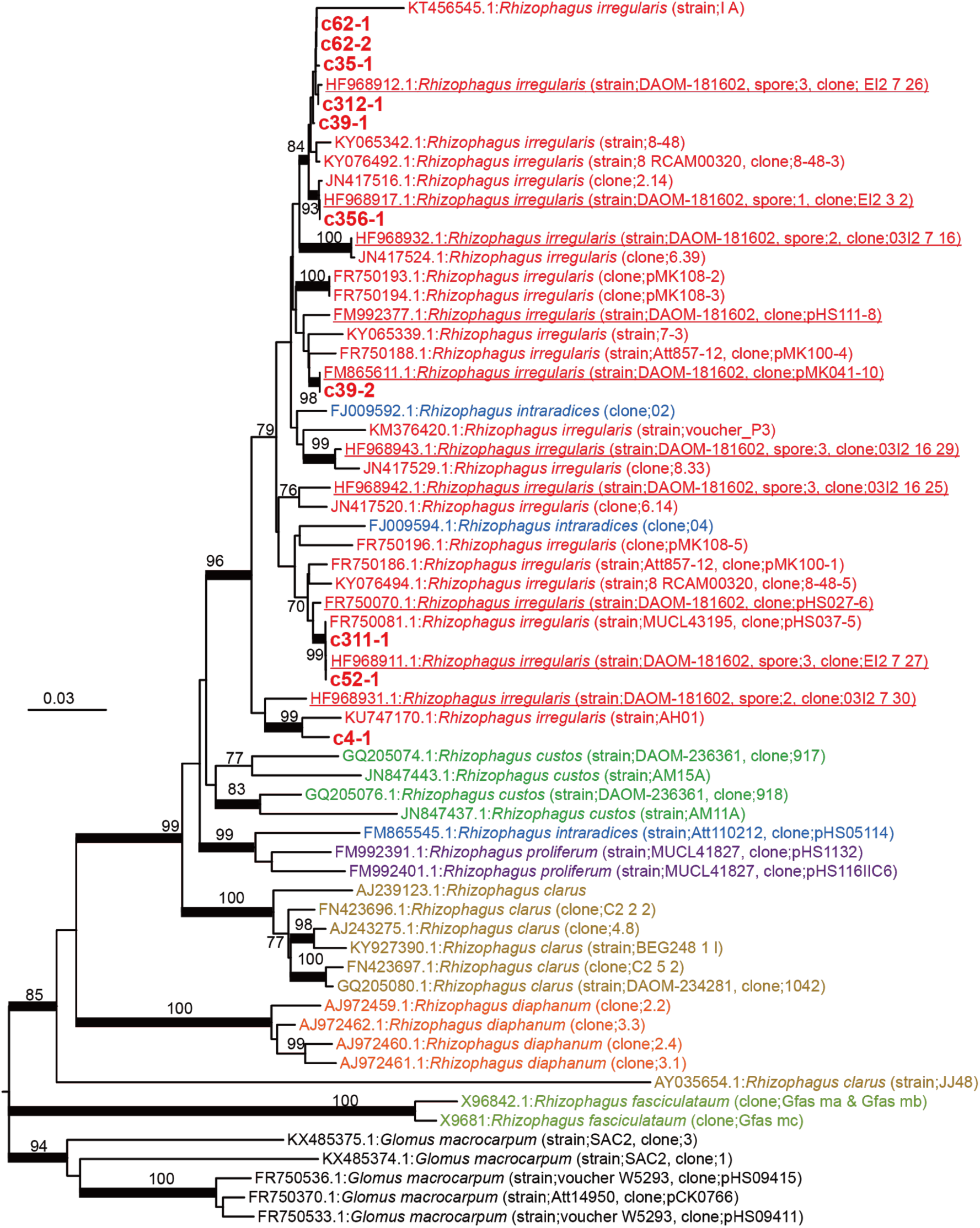
NJ tree based on 586 positions of 48S rDNA. Partial 18S, ITS1, 5.8S, ITS2 and partial 28S rDNAs were used. The ten rDNA paralogs from RIR17 and 58 *Rhizophagus* sequences from the DDBJ were chosen as operational taxonomic units (OTUs). The 58 *Rhizophagus* sequences were selected from 329 OTUs in the DDBJ (209 OTUs for DAOM-181602, 57 OTUs for other *R. irregularis* strains, and 63 OTUs for other *Rhizophagus* species) using CD-Hit clustering (-c 0.98 -n 5). Five *Glomus* sequences were used as outgroup OTUs. Red underlined OTUs are sequences from *R. irregularis* DAOM-181602, and other red OTUs are data from other strains of *R. irregularis*. Nodes supported by over 80 bootstrap values are marked by a bold line. All *R. irregularis* OTUs made a single clade with *Rhizophagus intraradices* that is a morphologically non-distinct sister group of *R. irregularis*.

**Table 2.** Numbers of intragenomic polymorphic sites in fungal rDNAs.

| Species | #<br>polymorphic<br>sites | Repeat unit<br>length (bp) | # of units<br>in genome | # of polymorphic<br>sites/100 bases |
| --- | --- | --- | --- | --- |
| *Rhizophagus irregularis* | 238 | 5847 | 10 | 4.07 |
| *Rhizophagus irregularis*<sup>13</sup> | 38 | 1563 | - | 2.43 |
| *Ashbya gossypii*<sup>30</sup> | 3 | 8147 | 50 | 0.04 |
| *Saccharomyces paradoxus*<sup>30</sup> | 13 | 9103 | 180 | 0.14 |
| *Saccharomyces cerevisiae*<sup>30</sup> | 4 | 9081 | 150 | 0.04 |
| *Aspergillus nidulans*<sup>30</sup> | 11 | 7651 | 45 | 0.14 |
| *Cryptococcus neoformans*<sup>30</sup> | 37 | 8082 | 55 | 0.46 |
| *Phoma exigua* var. *exigua*<sup>86</sup> | 27 | 1672 | - | 1.61 |
| *Mycosphaerella punctiformis*<sup>86</sup> | 26 | 1669 | - | 1.56 |
| *Teratosphaeria microspora*<sup>86</sup> | 16 | 1671 | - | 0.96 |
| *Davidiella tassiana*<sup>86</sup> | 33 | 1672 | - | 1.97 |

### A model for the relaxation of rDNA homogeneity in *R. irregularis*

The revealed non-tandem structure of AMF rDNA led to a model for the mechanism responsible for its intragenomic heterogeneity. Pawlowska and Taylor (2004) predicted that rDNA heterogeneity is caused by relaxation of “concerted rDNA evolution” in Glomerales including *Rhizophagus*^25^. However, details of the “relaxation” have been unclear. Here, we propose a hypothetical mechanism: the loss of TRSs precludes the presence of DNA conformations associated with rDNA amplification and the maintenance of its homogeneity. The standard model of “concerted rDNA evolution” needs two or more tandemly repeated rDNA segments because the rDNA duplicates using tandemly repeated rDNAs as binding sites and templates for replication (Supplementary Fig. 1c)^52^. Although non-tandem rDNAs are rare in eukaryotes, this trend of heterogeneity in non-tandem rDNAs has been detected by laboratory systems as well as in wild organisms; *Arabidopsis thaliana* has one pseudogenic rDNA (lacking 270 bases of an important helix as rRNA) besides the main tandem repeat rDNA arrays^53, 54^, and the lack of rDNA tandem repeats in malaria-causing *Plasmodium* parasites^49, 55^ indicates intragenomic rDNA polymorphisms. These observations support our hypothesis that rDNA heterogeneity in AMF is related to their lack of TRSs. AMF species may not amplify their rDNA by the general eukaryotic rDNA amplification system (USCR), which may increase their rDNA heterogeneity.

On the other hand, our phylogenetic analysis suggests that AMF has a system to maintain weak similarity among the paralogs without TRSs. Previously observed rDNA heterogeneity in Glomerales suggests that concerted evolution was relaxed before the diversification of *Rhizophagus* species^25, 29^. When the observed ten rDNAs duplicated before speciation and evolved independently, each of the duplicated genes formed a clade with orthologs in other species. However, we found no orthologous rDNA genes from other *Rhizophagus* species (Fig. 5). Our tree suggests that the observed rDNAs in *R. irregularis* either expanded or were assimilated after speciation. One hypothetical mechanism that would cause this similarity is homologous recombination via “synthesis-dependent strand annealing” (Supplementary Fig. 6)^56^. This conserved system to repair double-strand breaks (DSBs) results in non-crossover recombination and gene conversion wherein nonreciprocal genetic transfer occurs between two homologous sequences (Supplementary Fig. 6). Decreases in divergence by gene conversion are widely observed in duplicated genes. RIR17 showed that two rDNA pairs on the same contigs (c39-1 and c39-2, c62-1 and c62-2) had higher similarity than other paralogs (Fig. 4c). This similarity may be caused by the high gene conversion rate between these loci.

Our model raises a new question about the mechanism that maintains the number of rDNAs without gene duplication by USCR. Even if rDNA lacks TRSs, crossover recombination and single-strand annealing delete paralogous genes. Observed inverted repeat structures between rDNAs in proximity may contribute to inhibiting “single-strand annealing” between them and prevent copy number reduction. Plastidial rDNAs of land plants also make inverted repeat structures and conserve two rDNA copies on their plastidial DNA. Another probable system is the suppression of crossover recombination by the limitations of meiosis. When Holliday junctions dissociate without crossover, DSBs are repaired without gene number reduction. The majority of these crossovers arise during meiosis in eukaryotes^56^, and sexual reproduction had never observed in AMF. AMF species may keep their rDNA copy number by asexual spore-making and the rarity of their meiotic cell division.

### RNA-level impact and probable biological significances of non-tandemly repeated rDNAs

To confirm the transcriptional activity of each rDNA, we conducted total-RNA-Seq (RNA sequencing without poly-A tail selection. See material & method section.). Illumina sequencing of a modified library for rRNA sequencing (“Rir_RNA_rRNA” in Supplementary Table 2) produced 18,889,290 reads (read length = 100-301 bases) from DAOM-181602. We mapped the reads to all gene models from RIR17 (43,675 protein-encoding isoforms and ten 48S rDNA paralogs) and estimated the expression levels of each gene by eXpress software^57^. All rDNA paralogs had over 5,000 FPKMs (Fragments Per Kilobase of exon per Million mapped fragments) (Table 3), and multiple reads were matched to the specific region of each paralog, indicating that the ten rDNA copies are transcriptionally active. In general, eukaryotes silence a part of the rDNA copies^58^, and some eukaryotes change the transcribed rRNA sequences by “RNA editing”^59^. These editing and silencing processes were not detected in the AMF, and the rRNA were as polymorphic as the rDNA. These results show that DAOM-181602 has multiple types of ribosomes, each containing different rRNAs. Additionally, we detected highly duplicated ribosomal protein genes (e.g., ribosomal protein S17/S11) (Supplementary Tables 6 and 21) and tRNA genes, indicating unknown amino acid isotypes, which may also account for the heterogeneity of ribosomes (Supplementary Table 22).

The evolutionary significance of the of non-tandemly repeated heterogeneous rDNAs is unclear. One of the probable factors is a reduction in the need to maintain numerous rDNAs in a genome. As described in the above sections, the AMF rDNA copy number suggests a system that efficiently produces rRNA from a few rDNAs, and the inverted repeats structure of rDNAs and asexual spore reproduction will also reduce the deletion rate of rDNAs. AMF may thus no longer need to rapidly amplify rDNA copies using TRSs, and the slowed replacement rate of rDNA may then cause the heterogeneity as a side effect. Another possibly significant effect is the enhancement of phenotypic plasticity by ribosomal heterogeneity (Fig. 6). Recent studies have started to reveal that various eukaryotes (e.g., yeast, mice, and *Arabidopsis*) produce heterogeneous ribosomes and subsequently alter phenotypes via proteins translated by particular ribosomes^60^. Accelerated accumulation of AMF rDNA mutations by the lack of TRSs may lead to functional variety in produced ribosomes and increases in the rate of adaptation by different translation activities within the same species. Although the functional effects of observed rDNA mutations remain to be determined, the middle area of our 28S rDNA (4,450-4,500 bases on c62-1) had a higher mutation rate than ITS regions (Fig. 4a). Because the ITS regions (encoding non-functional RNA) vary under neutral mutation rates, the accumulated variants in the middle-28S region may have functional effects favored by natural selection (via diversifying selection). This region is thus a useful target for the future functional analyses of AMF rRNA.

**Fig. 6.**
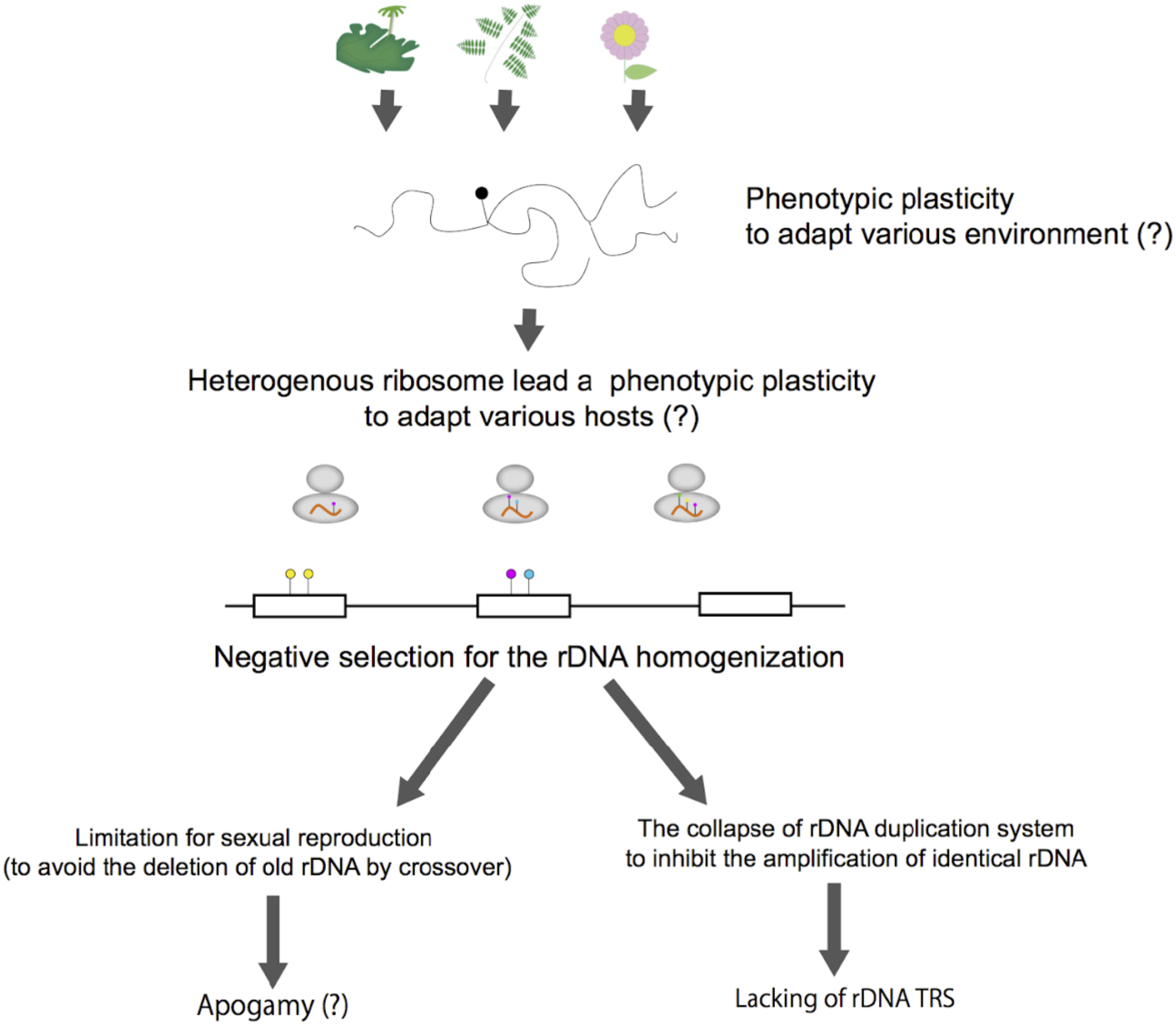
Hypothetical model for the evolution of unique rDNAs/rRNAs in AMF. Evolutionary model for the lack of TRSs in AMF and its sequence heterogeneity. The various environmental conditions (e.g., various host species) may lead to the evolution of phenotypic plasticity via multiple types of ribosomes in AMF. If the rDNA is exposed to disruptive selection, rDNA duplication by TRSs and USCR may be nonadaptive because the duplication of particular rDNA types reduces the variety of rDNA types. Sexual reproduction, combined with crossover recombination, may also be limited to inhibit the reduction of mutated rDNA.

AMF species are similar to the malaria parasite in that they both have heterogeneous non-tandem rDNAs and infect distantly related host species^49^. In the malaria parasite, changes in the ribosome properties depend on the host (human or mosquito), which is likely able to alter the rate of translation, either globally or of specific messenger RNAs, thereby changing the rate of cell growth or altering patterns of cell development^49^. The relationship between the diversity of host organisms and rDNA polymorphisms will be an important area for further research. The phenotypic plasticity caused by heterogeneous translation machinery may allow adaptation for various host species having slightly different symbiotic systems. Previous studies have proposed that the heterokaryosity in AMF species drives variable genetic combinations of mycelia in the absence of sexual recombination^61^. Recent genomic studies, furthermore, discovered signatures of sexual reproduction within the dikaryon-like stage^16, 62^. Our hypothesis does not exclude current theories for the genetic and phenotypic plasticity of AMF species (heterokaryosis and sexual reproduction) but proposes a multilayered diversification mechanism leading to their widespread distribution.

## Materials and Methods

### PacBio-based assembling

#### DNA preparation

The DNA sample for the PacBio and Illumina sequencing was extracted from a commercial strain of *R. irregularis* DAOM-181602 (MYCORISE® ASP, Premier Tech Biotechnologies, Canada). The DNA extraction followed the method of Fulton et al., 1995^63^ with some modifications described below. Purchased spore suspensions (including approximately 1,000,000 spores) were centrifuged (4500 rpm, 20 min), and washed three times with distilled water. Precipitated spores were frozen with liquid nitrogen, ground with pestle, and dispersed in extraction buffer (100 mM Tris-HCl pH 8.0, 20 mM EDTA, 0.75% sarkosyl, 0.1% PVP, 0.75% cetyl trimethylammonium bromide (CTAB), 0.13 M sorbitol, 0.75 M NaCl, and 0.1 mg/ml proteinase K). After incubation at 37 °C for 10 min, the aqueous phase was centrifuged (15000 rpm, 4 min), and the pellet was discarded. An equal volume of phenol/chloroform (1:1, vol/vol) was added, and the sample was gently mixed and centrifuged (15000 rpm, 2 min). The aqueous phase was collected, and an equal volume of chloroform was added to the sample, which was then mixed and centrifuged (15000 rpm, 2 min). The aqueous phase was collected again, and 1/10 vol of sodium acetate and 0.7 vol of isopropanol were added. The sample was then mixed and centrifuged (12000 rpm, 20 min). The resulting pellet was washed twice with 70% EtOH and eluted with TE buffer. Extracted DNA was purified with Genomic-tip (Qiagen, Germany) following the manufacturer’s instructions.

#### PacBio sequencing

Long-read sequences were generated with a PacBio RS II sequencer (Pacific Biosciences, Menlo Park, CA, USA) using a DNA/Polymerase Binding Kit P6 v2 (Pacific Biosciences) and a DNA Sequencing Reagent Kit 4.0 (Pacific Biosciences). The library was prepared according to the 20-kb Template Preparation Using BluePippin™ Size-Selection System (Sage Science, MA, USA). A total sequence of 11.7 Gb in 955,841 reads (76× coverage of the genome, assuming a genome size of 154 Mb) was obtained from 29 SMRT cells (Supplementary Table 2). The N50 length of the raw reads was 13,107 bases.

#### PacBio-based genome assembly

The *R. irregularis* genome was assembled using the RS_HGAP<u>_</u>Assembly.3 protocol for assembly and Quiver for genome polishing in SMRT Analysis v2.3.0 (Pacific Biosciences). The procedure consisted of three parts, involving (1) generation of preassembled reads with improved consensus accuracy; (2) assembly of the genome through overlap consensus accuracy using Celera Assembler; and (3) one round of genome polishing with Quiver. For HGAP, the following parameters were used: PreAssembler Filter v1 (minimum subread length = 500 bases, minimum polymerase read quality = 0.80, minimum polymerase read length = 100 bases); PreAssembler v2 (minimum seed length = 6,000 bases, number of seed read chunks = 6, alignment candidates per chunk = 10, total alignment candidates = 24, minimum coverage for correction = 6, and BLASR options = ‘noSplitSubreads, minReadLength = 200, maxScore = 1,000, and maxLCPLength = 16’); AssembleUnitig v1 (genome size = 150 Mb, target coverage = 25, overlapper error rate = 0.06, overlapper min length = 40 bases and overlapper k-mer = 14); and BLASR v1 mapping of reads for genome polishing with Quiver (maximum divergence = 30, minimum anchor size = 12). Assembly polishing with PacBio reads was carried out with Quiver using only unambiguously mapped reads. The statistics of the PacBio-only assembly set and previously sequenced data (Lin14, JGI_v1.0, JGI_v2.0) were evaluated using QUAST ver. 4.3^64^. The percentage of genome coverage was estimated assuming the genome size to be 154 Mb based on Tisserant et al^17^.

#### Error correction with HiSeq data and identification of host plant contamination

After polishing using Illumina data, we eliminated the sequences derived from contaminated DNAs during the sample preparation. BLASTn search of the polished assemblies against the “refseq_genomic” database detected nine assemblies showing similarity with sequences from carrot (BLAST ver. 2.2.31+, query coverage per subject >95%, percentages of identical matches >90%, bit score > 1000) (Supplementary Table 2), which might be used as a host plant by the manufacturer for the cultivation of *R. irregularis* samples. After elimination of the nine contaminated contigs, we submitted the assemblies to the DDBJ as whole-genome shotgun sequence data (RIR17) of *R. irregularis* DAOM-181602 (BDIQ01).

#### Genomic alignment with previous genome assemblies

The quality of our genome assembly was evaluated by alignment with previously available *R. irregularis* DAOM-181602 genome assemblies. A one-by-one genome alignment was constructed by MUMmer ver. 4.0.0beta2^65^ between RIR17, JGI_v2.0, Lin14, and JGI_v1.0 assemblies. Each genome set was aligned by the nucmer function in MUMmer, and the statistics of the alignments were extracted by the dnadiff wrapper with the default setting.

### Gene prediction and annotation

*De novo* repeat motifs were identified using RepeatModeler ver. 1.0.8, which combines RECON and RepeatScout programs^37^. Based on the identified motif, the repetitive region in the assemblies was masked with RepeatMasker ver. 4.0.5^37^. We used the default parameters for the identification and the masking.

For the gene models constructed from RIR17 assemblies, standard RNA-Seq data were obtained from *R. irregularis* spores and hyphae. The RNA was extracted with an RNeasy Plant Mini kit (Qiagen) after incubation of the purchased spores (MYCORISE® ASP) in a minimum nutrient medium for one day. An Illumina RNA-Seq library was constructed with a TruSeq Stranded mRNA Library prep kit (Illumina). The library was sequenced (101 bases from each end) on a HiSeq 1500 platform (Illumina). A total of 16,122,964 raw reads (3.2 Gb) were obtained from the library (Supplementary Table 2). After filtering low-quality and adapter sequences, RNA-Seq data were mapped to RIR17 assemblies with TopHat ver. 2.1.1^66^ with the default setting.

Then, the RIR17 assemblies were processed through the RNA-Seq-based gene model construction pipeline using AUGUSTUS ver. 3.2.1 software^67^. We constructed *R. irregularis*-specific probabilistic models of the gene structure based on 495 manually constructed gene models from the longest “unitig_392” sequence in RIR17. Manual gene models were made with *ab initio* AUGUSTUS analysis based on probabilistic models for *Rhizopus oryzae* and by manual refinement using the homology data with already-known genes and mapped RNA-Seq data. Then, with the trained probabilistic models and the intron-hints data from the mapped RNA-Seq read, 37,639 optimal gene models were constructed using the AUGUSTUS pipeline. We then confirmed whether the AUGUSTUS pipeline overlooked the called genes in previous genome studies. We mapped all transcript sequences obtained from previous gene modeling on Lin14 and JGI_v1.0 against our RIR17 genomic sequences with Exonerate^68^ (ver. 2.2.0, option --model est2genome --bestn 1), resulting in the recruitment of 3,933 overlooked genes. The completeness of the constructed gene model was evaluated with BUSCO ver. 2.0^39^. The BUSCO analysis used the “Fungi odb9” gene set (http://buscodev.ezlab.org/datasets/fungiodb9.tar.gz) as a benchmark and employed the “-m proteins” option to analyze the preconstructed protein data without the *ab initio* gene modeling step.

The confidences of the obtained 41,572 gene models were estimated based on 1) RNA-Seq expression support, 2) homology evidence, and 3) protein motif evidence. For the calculation of gene expression levels, we mapped our “Rir_RNA_SS” data and 32 RNA-Seq data submitted to the sequence read archive (SRA) database (24 data sets from DRP002784 and 8 data sets from DRP003319) and calculated the gene expression levels (FPKM) using FeatureCounts^69^ with the default setting (Supplementary Table 6). Homology with previously known genes was determined by BLAST searches against the orthoDB (odb9) (Supplementary Tables 6 and 21). The protein motif was searched using Pfam analysis in InterProScan ver. 5.23-62.0^70^ (Supplementary Table 6).

Constructed gene models were annotated by several *in-silico* searches. Gene functions were predicted based on BLASTp (Database = nr, RefSeq and UniProt), and Pfam in InterProScan (Supplementary Table 6). We manually selected the descriptive nomenclatures from those four searches and submitted to the DDBJ. Orthologous relationships were classified with Orthofinder (ver. 1.1.2)^71^, and rapidity expanded/contracted families were analyzed with CAFE (ver. 4.1)^72^ from Orthofinder results. Phylogenetic trees for the CAFE analysis were constructed with IQ-tree (ver. 1.6.1)^73^ for maximum likelihood (ML) analysis and r8s (v1.81) for a conversion for an ultrametric tree. An ML tree was made from 159 single-copy genes from the Orthofinder results (Supplementary Table 6) and was converted to an ultrametric tree based on the divergence times of AMF-Mortierellales (460 Myr)^27^ and Deuterostomia-Protostomia (550 Myr)^74^. Overlapping genes with TEs were extracted from AUGUSTUS and RepeatMasker results using bedtools (ver. 2.26.0, “bedtools intersect” with -wa option)^75^.

The MACG (missing ascomycete core gene) orthologs were sought using BLAST with the “-evalue 0.0001” option, and the reference sequences for the MACG search were selected from protein data from an S288C reference in the *Saccharomyces* genome data base (SGD) (Supplementary Table 15). Genes involved in the degradation of plant cell walls were sought by BLAST with the same settings as the MACG search, and the reference sequences were selected from *Aspergillus niger* CBS 513.88 data in GenBank based on CAZY classification (Supplementary Table 15). Other gene annotations based on the CAZy database were performed with the dbCAN HMMs 6.0 web service^76^ (Supplementary Table 6).

### Detection of Ribosomal DNA and intragenomic polymorphisms

Ribosomal DNA regions were detected by RNAmmer ver. 1.2^77^ from whole RIR17 assemblies and were manually refined based on the MAFFT v7.294b^78^ alignment to the 48S rRNA in *Saccharomyces cerevisiae* S288C. The genomic positions of rDNAs were visualized with Python ver. 3.4.0 (BasicChromosome ver. 1.68, and GenomeDiagram ver. 0.2 modules) (Fig. 2a).

The number of rDNA paralogs in the genome was estimated by mean depth of coverage. We masked repetitive regions (based on RepeatModeler analysis) and all rDNA regions on RIR17 except one rDNA copy (c62-1). Then, trimmed R1 Illumina reads from “Rir_DNA_PE180” library were mapped to the repeat-masked RIR17 using bowtie2 ver. 2.2.9^79^. The coverage depth of the rDNA region and 243 single-copy BUSCOs were obtained using bedtools (“bedtools coverage” command with -d option), and the statistics of each region were calculated and visualized by R software ver. 3.4.2 with the ggplot2 library (Fig. 2b). To prevent copy number estimation from depth fluctuation due to the intragenomic heterogeneity, we confirmed the coverage depth using the consensus sequences of all ten rDNA paralogs; the joined Illumina reads (from “Rir_DNA_PE180” library) were mapped back to a consensus rDNA sequences and ten single-copy BUSCO genes from RIR17, and the depth of coverages was then counted by bedtools (genomeCoverageBed) (Supplementary Table 17).

The syntenic structure around rDNA genes was confirmed by the mapping of PacBio raw reads and comparison with JGI_v2.0 assemblies. All of the “filtered-subreads” from SMART Analysis software were mapped to RIR17 assemblies by BWA-MEM (ver. 0.7.15-r1140) with the “-x pacbio” option. Mapped reads were visualized with Integrative Genomics Viewer (ver. 2.4), and the reads covering the rDNA regions were selected by eye. Alignment between JGI_v2.0 and RIR17 was done by a combination of MUMmer, LASTz (ver. 1.04.00), and AliTV^80^ (ver. 1.0.4) software. JGI_v2.0 scaffolds having regions corresponding with RIR17 sequences were selected by the nucmer and delta-filter (with -1 option) functions in MUMmer. Then, we extracted the JGI_v2.0 scaffolds corresponding to RIR17 contigs with rDNAs (“unitig_311”, “_312”, “_35”, “_356”, “_4”, and “_52”). Selected scaffolds were aligned to the corresponding RIR17 contigs by alitv.pl scripts (with “alignment: program: lastz” and “--ambiguous=n” settings) and alitv-filter.pl (with “--min-link-identity 80” and “--min-link-length 10000” option) in the AliTV package and visualized with the AliTV web service (http://alitvteam.github.io/AliTV/d3/AliTV.html).

The difference among the rDNA paralogs was calculated from the aligned sequences by MAFFT ver. 7.309 (options: --localpair, --op 5, --ep 3, --maxiterate 1000) using the pairwise comparison with CLC Main Workbench 7.8.1 (Qiagen). The mutation type was called by eye from the alignment, and we chose the c62-1 paralog as a reference sequence for mutation calling (Fig. 4a). Phylogenetic trees (Figs. 4c and 5) were constructed from the MAFFT alignment by the neighbor-joining method with MEGA^81^ ver. 7.0.21 under the maximum composite likelihood model and were tested for robustness by bootstrapping (500 pseudoreplicates).

### Heterogeneity of translation machineries

The expression levels of the rDNA paralogs were examined with modified Illumina sequencing of *R. irregularis* spores and hyphae. Total RNA was extracted with an RNeasy Plant Mini kit (Qiagen) after the incubation of the MYCORISE® spores in a minimum nutrient medium for seven days. An Illumina RNA-Seq library was constructed with a TruSeq Stranded mRNA Library prep kit (Illumina). To skip the poly-A tailing selection step in the library construction, we started from the “fragmentation step” of the standard manufacturer’s instructions. The library was sequenced (301 bases from each end) on a MiSeq platform (Illumina). A total of 16,122,964 raw reads (3.2 Gb) were obtained from the library (Supplementary Table 2). After filtering low-quality and adapter sequences, RNA-seq data were mapped to the RIR17 assembly with TopHat with the default settings. Fragments Per Kilobase of exon per Million mapped fragments (FPKMs) for each gene were calculated with eXpress ver. 1.5.1 with the “--no-bias-correct” option. Transfer RNAs were identified with tRNAscan-SE^82^ ver. 1.3.1.

## Data Availability

Raw reads, genome assemblies, and annotations were deposited at INSDC under the accessions as follows; Sequence read archive: DRA004849, DRA004878, DRA004889, DRA004835, DRA005204, and DRA006039; Whole genome assembly: BDIQ01000001-BDIQ01000210; Annotations: GBC10881-GBC54553. All the other data generated or analyzed during this study are included in this published article and its Supplementary information.

## Author contributions

T.M., S.S., M.K., conceived of and designed the experiments; T.M., Y.K., H.K., N.T., K.Y., and

T.B. performed the experiments; T.M., N.O., S.S., and T.B. analyzed the data, T.M., Y.K., H.K., K.Y., T.B., S.S., and M.K. wrote the manuscript.

## Acknowledgements

This work was supported by JST ACCEL Grant Number JPMJAC1403, Japan. We thank the “Functional Genomics Facility” and the “Data Integration and Analysis Facility” at the National Institute for Basic Biology for technical support; Katsuharu Saito, Kohki Akiyama, and two anonymous reviewers for discussions; and present and past members of the Kawaguchi lab and the Shigenobu lab.

---

**Supplementary Fig. 1.**
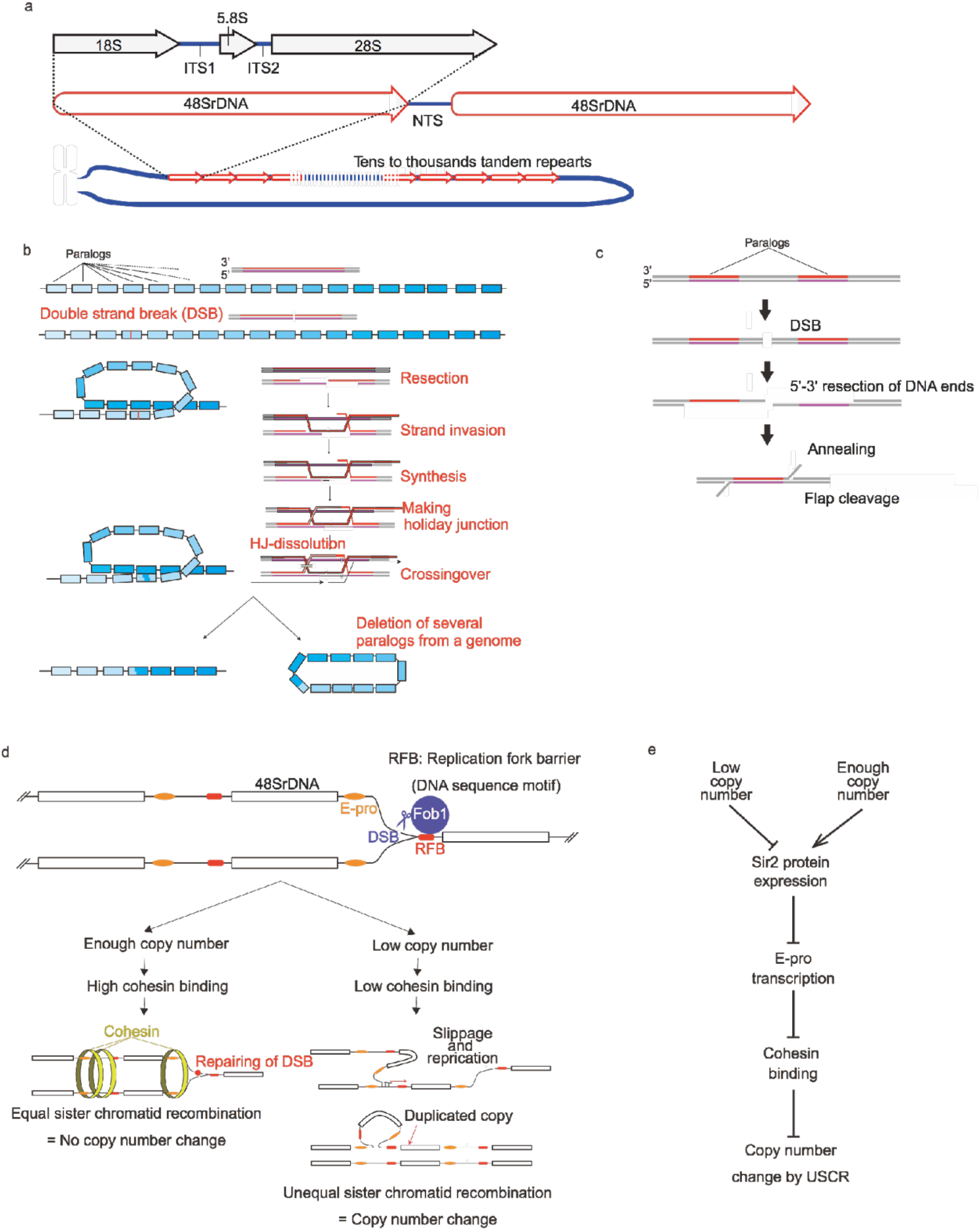
General concerted evolution of eukaryotic rDNA. a. A general structure of eukaryotic rDNA clusters^20^. b. Deletion of homologous genes by crossover recombination^33, 56^. c. Deletion of homologous genes by single-strand annealing^34^. d. A model of the rDNA number maintenance system^32^. e. Copy number-controlling pathway in yeast^32^.

**Supplementary Fig. 2.**
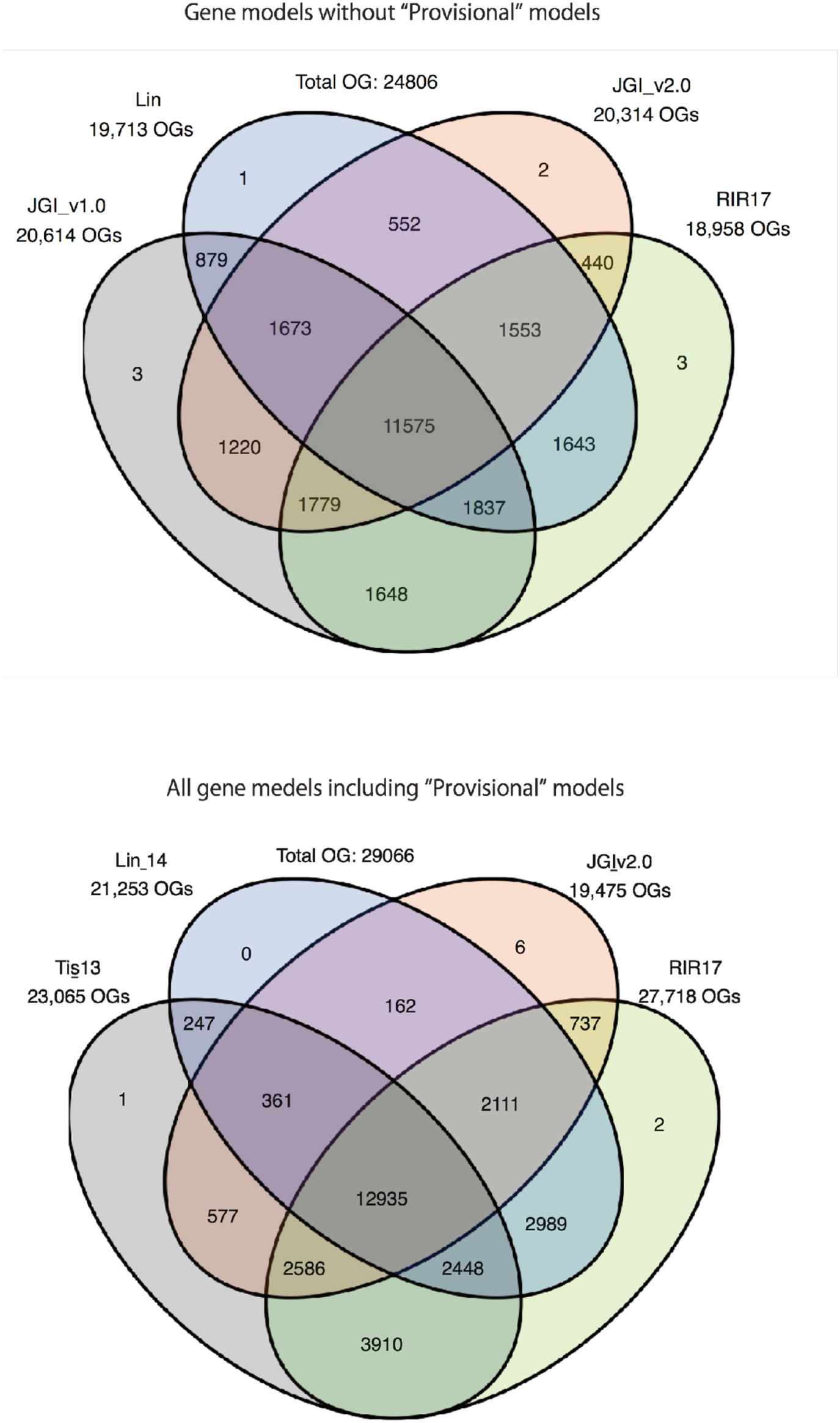
Cross comparison of *R. irregularis* DAOM-18160219 orthologous genes from three genomic studies and our RIR17 assemblies.

**Supplementary Fig. 3.**
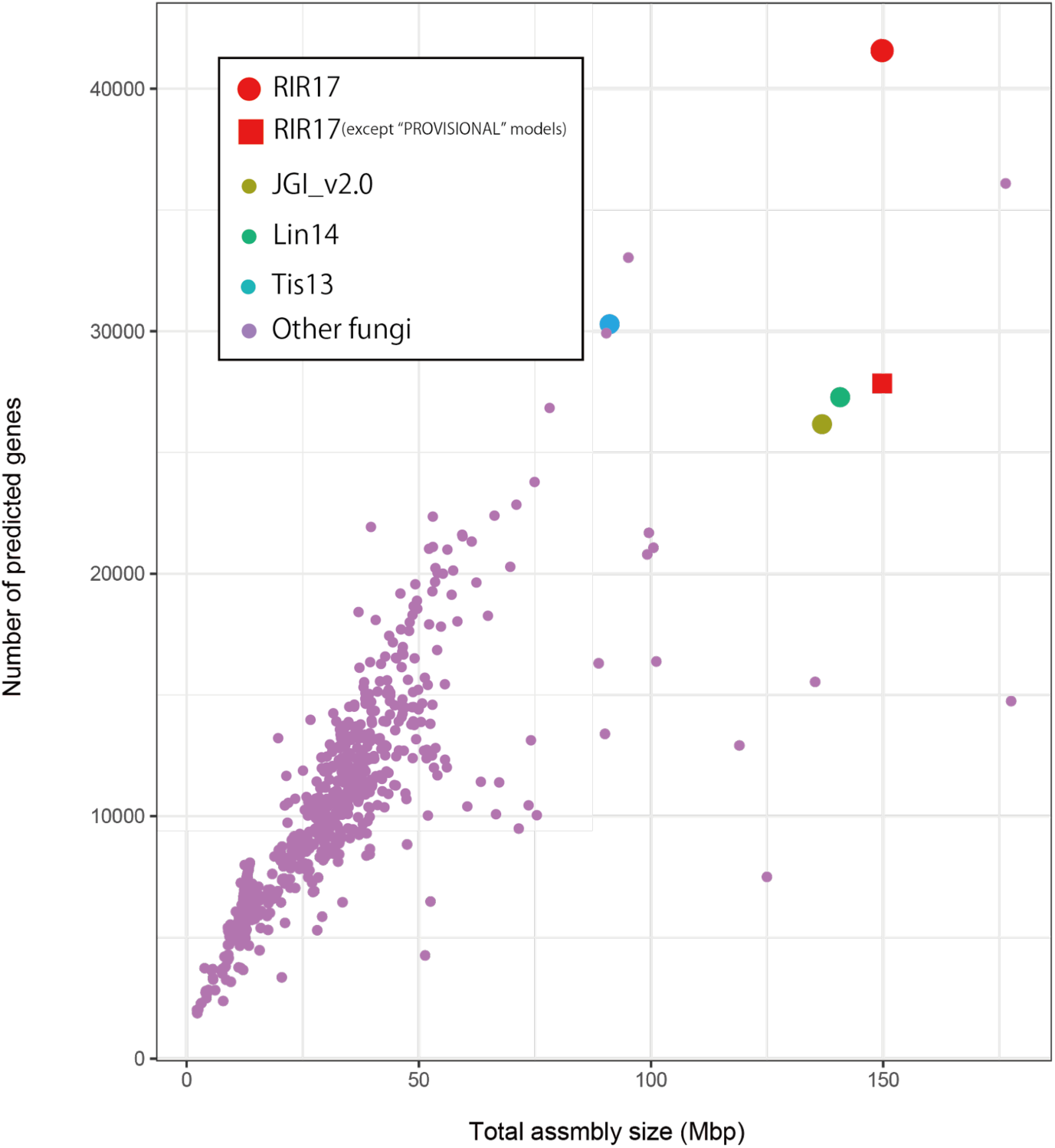
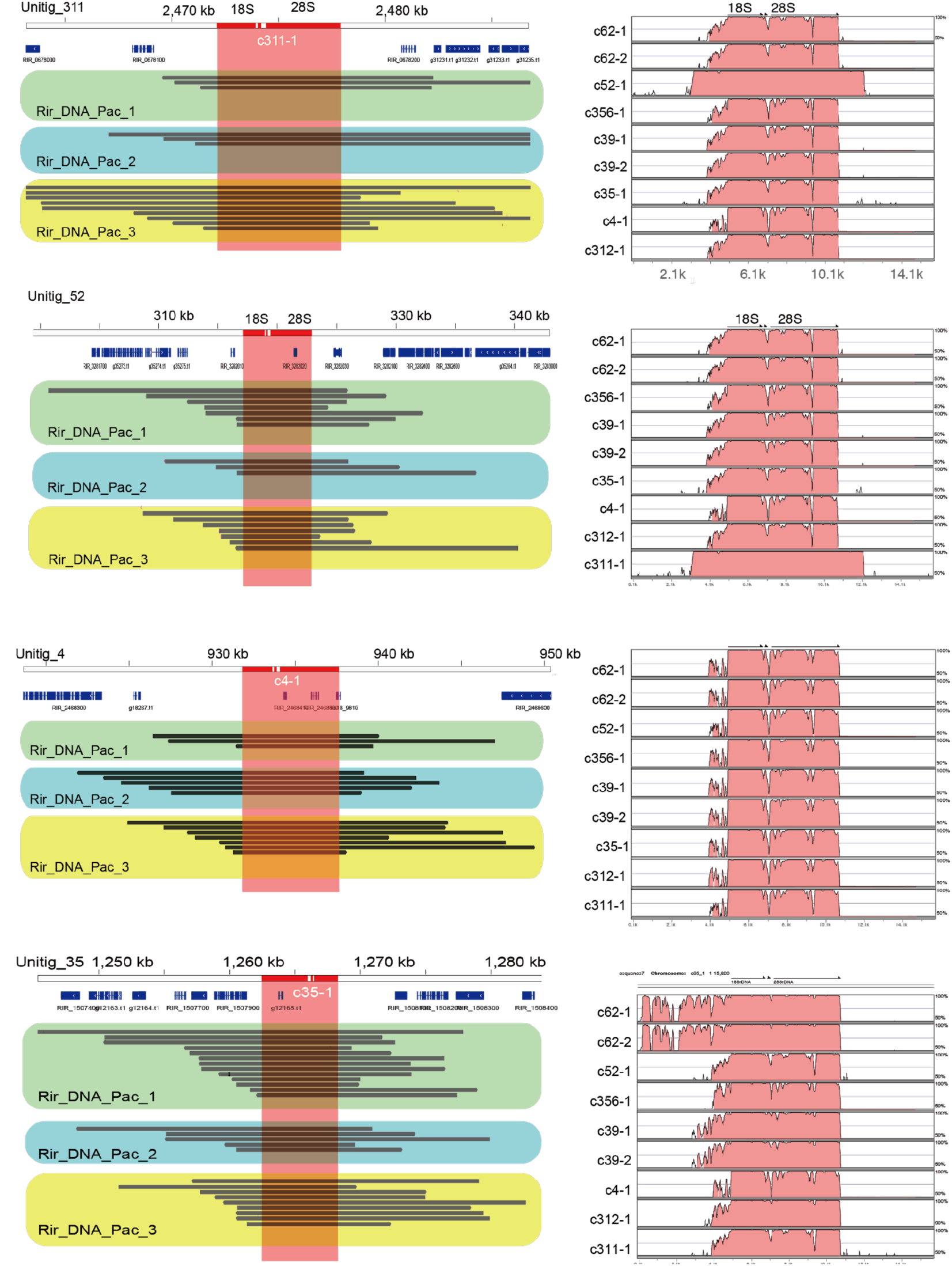
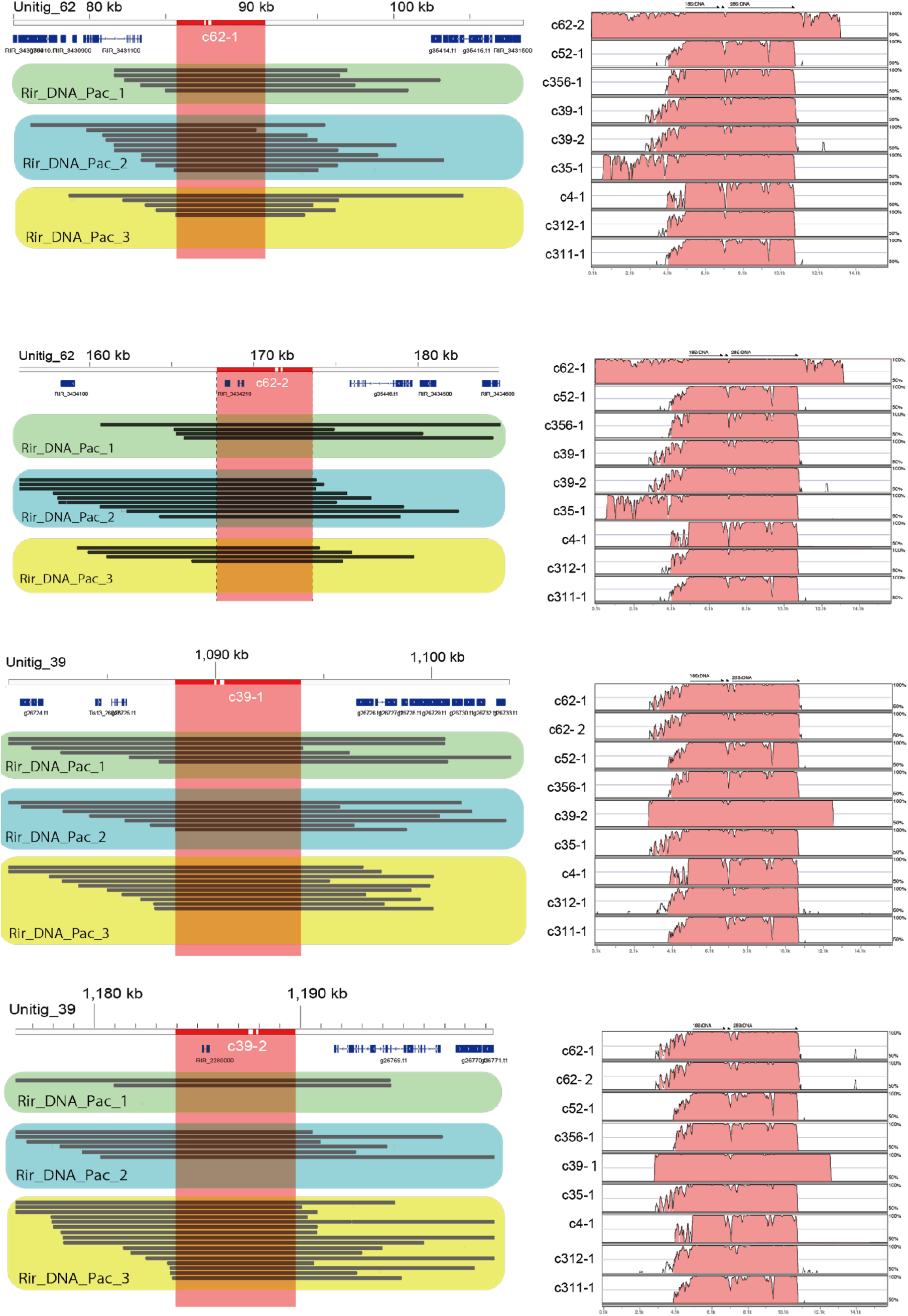
Total assembly size and predicted gene number in fungi. *Rhizophagus irregularis* genomes (RIR17 (this work) and two previously assembled genomes, JGI_v1.0 and Lin14) and 768 genomes registered in GenBank. The fungal assembly statistics were obtained from the registered information in GenBank (ftp://ftp.ncbi.nlm.nih.gov/genomes/ASSEMBLY_REPORTS/All/).

**Supplementary Fig. 4.**
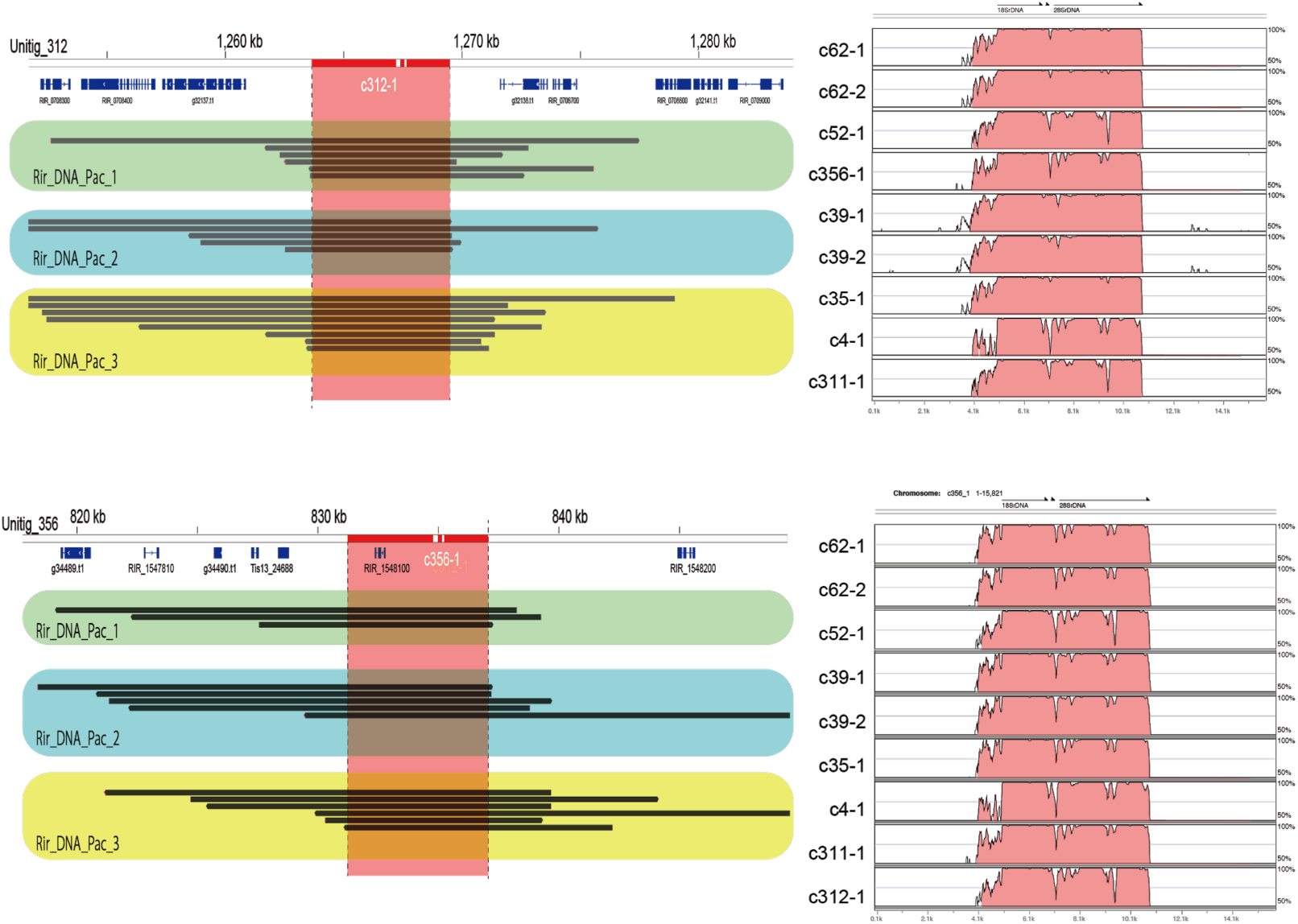
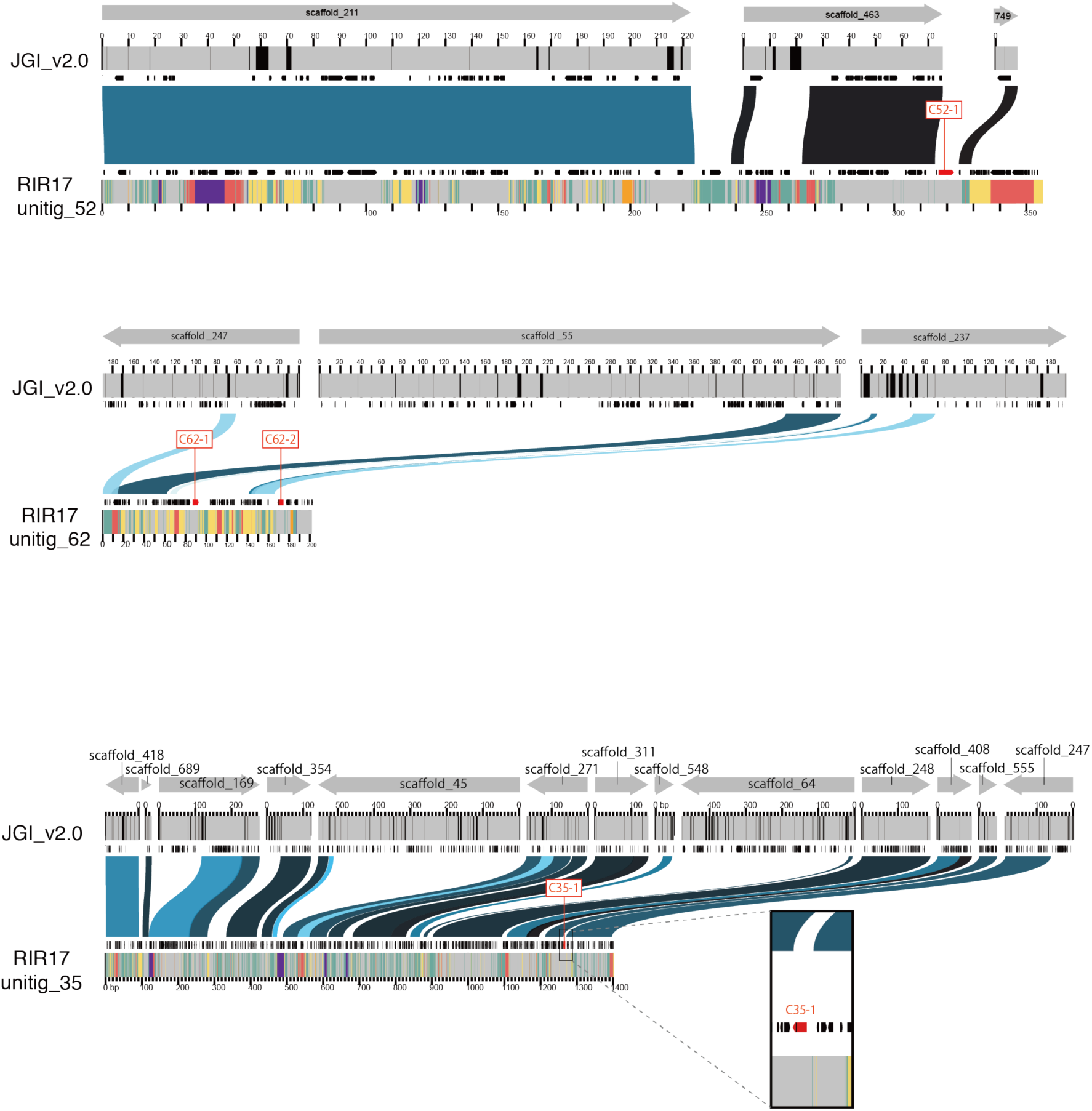
Mapped PacBio reads of rDNA regions of RIR17 contigs and the sequence similarity of rDNAs with other rDNA regions in RIR17 The colors have the same meanings as in Fig. 3a-b.

**Figure S5.**
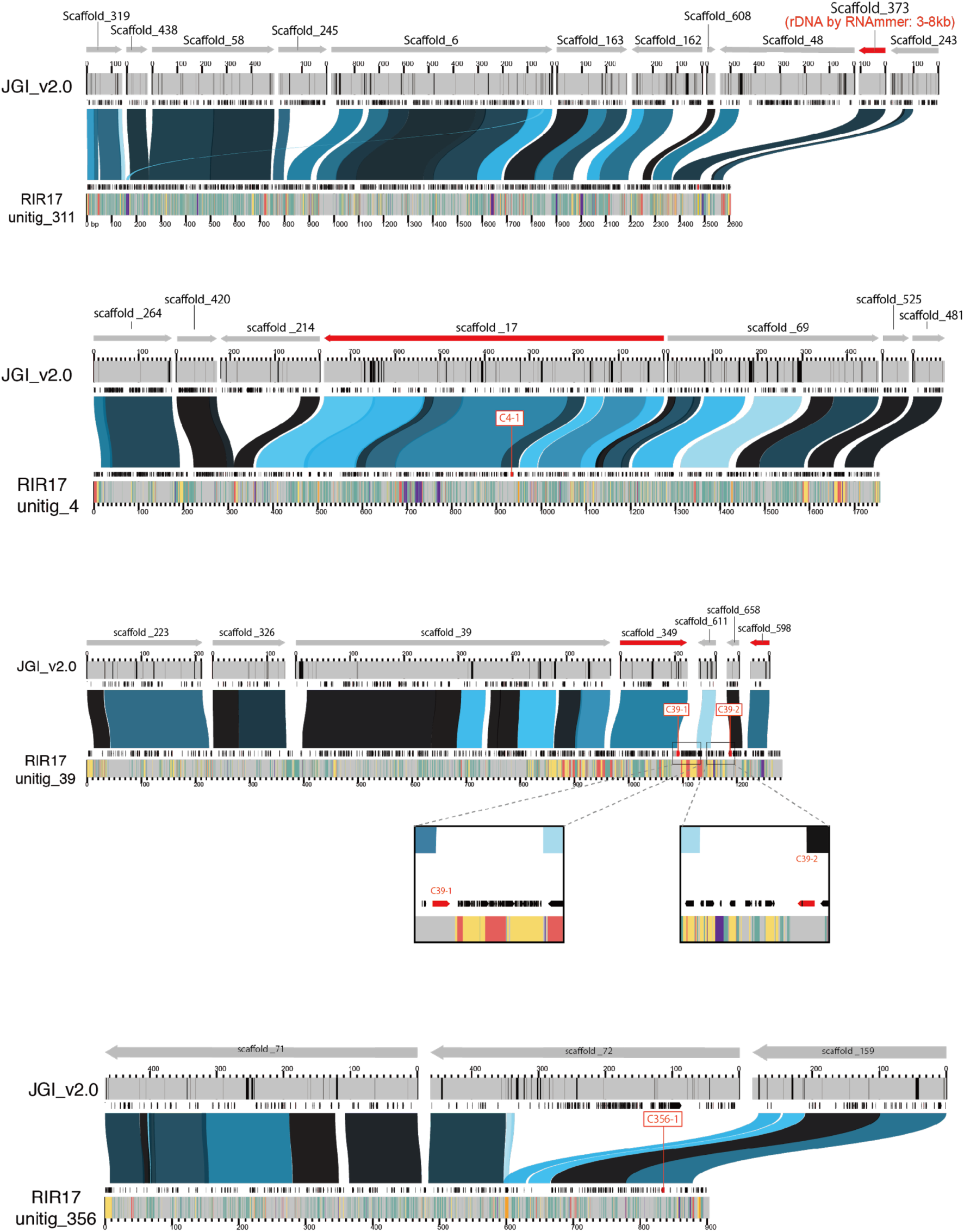
Positions and identities of JGI_v2.0 scaffold aligned against RIR17 contigs with rDNAs. The colors have the same meanings as in Figure 3c.

